# Generative AI Models Reveal Dynamic Views of Aging (DyViA) Phenotypes in Healthy Individuals

**DOI:** 10.64898/2026.07.05.735302

**Authors:** Deep Ray, Meghana Ray, Saumyadipta Pyne

**Author notes:** Corresponding author: Saumyadipta Pyne (SP). Emails: (DR); (MR).

## Abstract

**Background and objectives:** In recent years, the need to develop analytical strategies for healthy aging has assumed great importance. In this study, we introduce DyViA, a generative artificial intelligence (genAI) platform that can construct personalized trajectories capable of predicting the plausible progression of selected phenotypes with advancing age.

**Research design and methods:** DyViA presents a suite of deep learning models covering two major GenAI approaches: DyViA-Diff, a new diffusion model; and DyViA-mGAN, an improved version of a recent Generative Adversarial Network model. It demonstrated the dynamic progression of femoral neck bone mineral density (BMD) using data from a longitudinal cohort study of women in the U.S. of age 65 years or above.

**Results:** Using very few initial measurements, DyViA generated individual-specific continuous trajectories of BMD, with a corresponding region of acceptable predictions, from 66 to 89 years. The results were subjected to rigorous quality-control and comparative analysis across multiple methods. While DyViA-Diff is the superior model with more coherent and accurate predictions, DyViA-mGAN allows for encoding population- and individual-level effects with a better control.

**Discussion and implications:** Given the prevalence of osteoporosis in the aging population, the main impact of DyViA’s genAI-driven contribution in the form of personalized, plausible models of BMD progression with age lies in the systematic yet rigorous transition from otherwise static models of inference about a clearly dynamic phenomenon to a continuous one. The foresight offered by DyViA’s outputs empowers an individual by conferring a certain degree of strategic preparedness in the course of aging.

## Background and objectives

In the light of general increase in lifespan among many populations around the world, it is the extension of individual healthspan that is currently the key goal in aging research (Olshansky, 2018). In this regard, a past article made two fundamental observations: (a) “health itself may be better considered a continuous variable that changes in a dynamic way throughout life” and (b) “[t]he health trajectory will be different in different individuals but will generally trend downward with age” (Kaeberlein, 2018). While these points may appear to be intuitive, hardly any models exist in the literature that can take as input a few static and discrete observations of an individual’s certain phenotypes relevant to aging and predict the person-specific dynamic healthspan in the form of continuous trajectories over time ranging from the present to years or decades into the future.

In various health-related phenomena, while trajectories are used for modeling of trends or detecting vulnerable clusters, these are generally constructed at the level of either populations or groups therein (Cawthon et al., 2020; Guo et al., 2024; Siggaard et al., 2020). Our aim in this study – to produce realistic Dynamic Views of Aging (DyViA) of a given individual – is fundamentally different. DyViA is a new generative artificial intelligence (genAI) platform that can construct personalized trajectories capable of predicting the plausible progression of selected phenotypes with advancing age. Its prediction is based on static phenotypic measurements that are usually available for an individual at a (very) few initial time-instances along with potential data on other covariates of the same phenomenon. In particular, DyViA presents a suite of new deep learning (DL) models, which is used here for demonstrating the *dynamic* progression of femoral neck bone mineral density (BMD) – the selected phenotype of interest – in healthy individuals as they continue to age past 65 years. Our selection of the phenotype is driven by the fact that as individuals age, BMD decreases, and therefore, osteoporosis becomes more prevalent in older adults (Li et al., 2017).

In recent years, aging research and its related fields have seen numerous applications of machine learning (ML), DL and genAI (Wilczok, 2025). These include deep aging clocks, biomarker discovery, generation of therapeutics, uses of multimodal, multiomic, and synthetic data, etc (Wilczok, 2025; Zhavoronkov et al., 2019). As a prominent example of genAI, the Generative Adversarial Networks (GAN), (Goodfellow et al., 2020) have been remarkably effective in modeling various aging-associated phenomena, e.g., Alzheimer’s disease progression (Jung et al., 2023), realistic age-driven changes in the heart (Campello et al., 2022), virtual training programs for elderly (Yu & Dang, 2025). Diffusion models present another emerging DL approach to genAI, e.g., brain age prediction (Zapaishchykova et al., 2024).

Traditional models, on the other hand, have found extensive and specialized applications in studies on BMD, often based on Dual-energy X-ray Absorptiometry (DXA) measurements. However, even the DL applications in this area have focused mostly on producing results that are characteristically static. Either these are used upstream in the data analytics pipeline such as for DXA image analysis (e.g., (Ammar et al., 2024; Leong et al., 2024)) or downstream typically for clinical diagnosis of osteoporosis (e.g., (Naeem et al., 2025)) or fracture prediction (e.g., (Nissinen et al., 2021)). Yet, studies of healthspan are not conceptually limited to these endpoints as they usually involve longitudinal data from healthy individuals, and produce insights into a dynamic phenomenon that, we think, is best captured by continuous trajectories. In contrast, FRAX and such calculators of long-term risk of fractures output an endpoint probability score that neither generates nor represents any continuous progression of the underlying phenotype (Carey et al., 2022).

Interestingly, some recent studies involving traditional approaches such as group-based growth mixture model (Guo et al., 2024) and linear mixed model (LMM) (van der Veere et al., 2024) have represented longitudinal aging data in the form of trajectories. In particular, a conditional GAN-based platform, DyViA-GAN, has not only produced dynamic, continuous, and personalized trajectories of BMD as an aging phenotype, it also outperformed multiple traditional models (Pyne et al., 2026). In this study, our aim is to introduce a further improved version of that model, DyViA-mGAN, as well as a new DL-based diffusion model, DyViA-Diff, for the same purpose. Together, this suite of models comprises a multi-option GenAI platform called DyViA to synthesize aging phenotype dynamics. We demonstrated it using longitudinal data from cohorts comprising healthy individuals in the U.S. of age 65 years and above. DyViA generates individual-specific plausible and continuous trajectories of femoral neck BMD values from 65 to 89 years along with confidence intervals. The following sections describe the details of these models, their respective results and a comparison thereof, the limitations, and future directions.

## Research design and methods

The goal is to predict personalized trajectories of selected phenotypes given available measurements of the phenotype at a few initial time instances and additional risk factors. We begin by introducing some notations needed to describe the proposed methodology. Let *ρ(t)* denote the value of the phenotype of interest at time *t*. We assume access to an individual’s phenotype measurements at *K* time points, which we collectively denote using the list *ρ*^*hist*^ = [*t*_1_, *ρ*(*t*_1_), *t*_2_, *ρ*(*t*_2_) …, *t*_*K*_, *ρ*(*t*_*K*_)]. Let *Q* be list of additional covariates (static or dynamic risk factors) available for this individual. We are then interested in predicting the possible value of the individual’s phenotype at a new (future) time *t*.

This task can be formulated in a mathematically rigorous manner as learning the conditional probability distribution *px*_|*y*_(*x*|*y*) of a random variable *X* given a measurement *y* of another random variable *y*. In the context of the present study, choose we *X* = *ρ(t)* and the composite conditioning variable as *y* = (*ρ*^*hist*^, *Q, t*). By allowing the time point *t* to vary, we learn the probability distribution of personalized continuous trajectories given the known measurements.

We consider two deep learning-based models to learn and sample the conditional distributions: (i) a conditional generative adversarial network (cGAN) and (ii) a conditional score-based diffusion model (cSDM). Both models are trained on a labeled dataset of the form D = {(*x*^(*i*)^, *y*^(*i*)^)} for *i* = 1, …, *N*. We note that the available phenotype measurement *ρ*^*hist*^ is typically scarce. Furthermore, for any two samples (*x*^(*i*)^, *y*^(*i*)^) and (*x*^(*j*)^, *y*^(*j*)^) in the dataset, the time points at which the phenotype values are known may not be the same, i.e., the dataset is temporally unstructured. The workflow of the proposed methods is shown in Figure 1, which depicts three phases: Secure data acquisition (Phase 1), training the generative models (Phase 2), and predicting plausible trajectories using the trained models (Phase 3).

**Figure 1:**
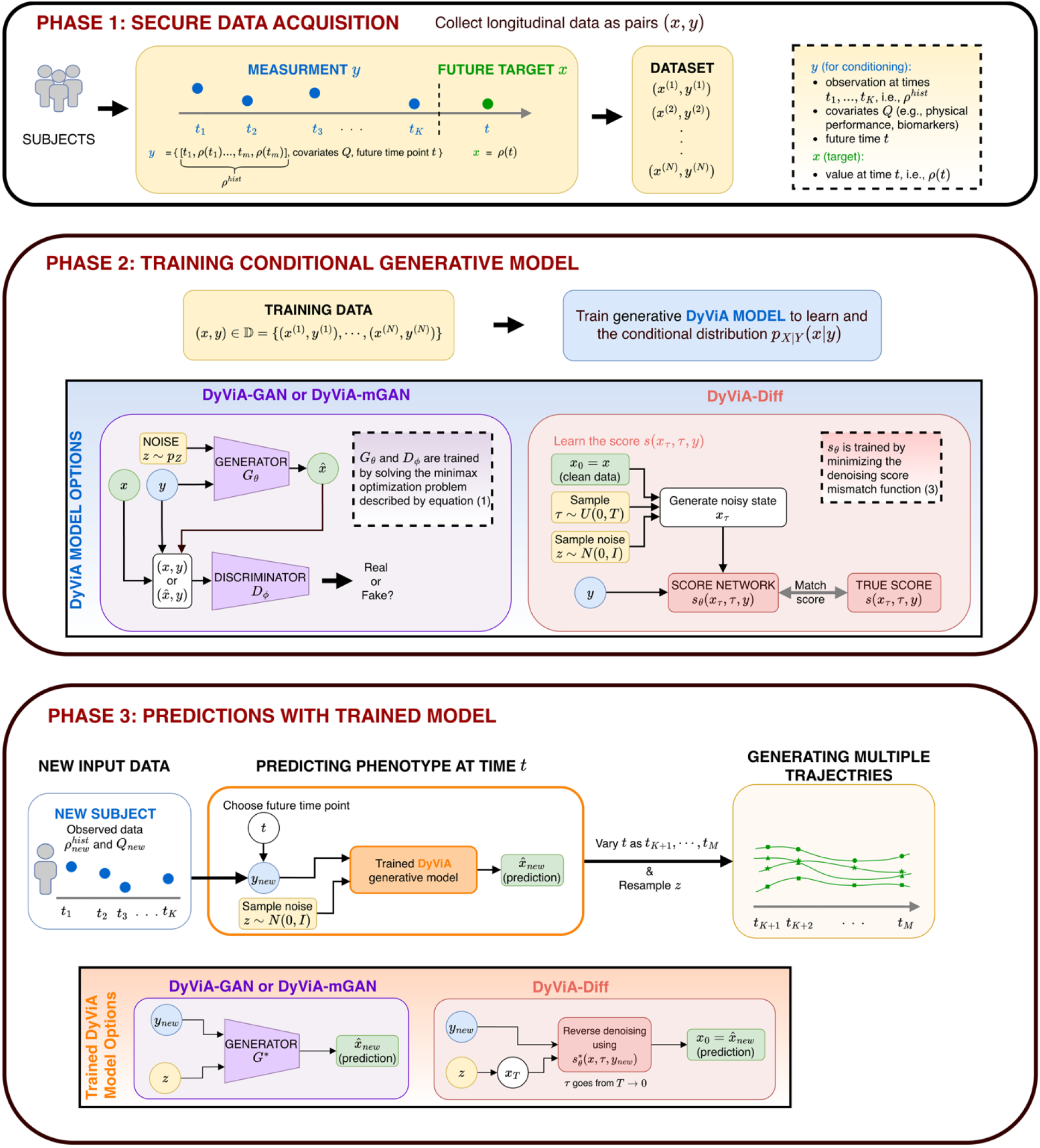
Schematic showcasing the process to design DyViA generative models. (Phase 1) Data acquisition for training and testing while following necessary security protocols. (Phase 2) Training the underlying neural networks describing the generative model. (Phase 3) Predictions for a new test subject conditioned on observed data y_new_. Predictions can be used to generate an ensemble of possible personalized trajectories by varying the future time point t and resampling the noise variable Z.

## DyViA-mGAN

Recently, a cGAN model called DyViA-GAN was proposed (Pyne et al., 2026) to learn personalized trajectories of bone density. We propose an improved variant called DyViA-mGAN which generates trajectories with mixed effects, i.e., models both population level and individual level dynamics.

DyViA-mGAN comprises two neural networks, namely the generator *G*_*θ*_ and the discriminator *D*_*ϕ*_ with trainable parameters *θ, ϕ* respectively. Given an input *y* = *y* (the measured data and the query time point *t*) and a latent variable *Z* sampled from a simple probability distribution *pz*, typically a standard Gaussian *N*(0, *I*)^1^ of size *N*_*Z*_, the generator constructs a close-to-real possible value of the phenotype at time *t*. On the other hand, the discriminator’s role is to distinguish between real values of the phenotype (available in the dataset) at time *t* and the fake values at the same time generated by *G*. The two networks are trained simultaneously in an adversarial fashion on the dataset D by solving a minimax optimization problem using the following adversarial loss (Mao et al., 2017)

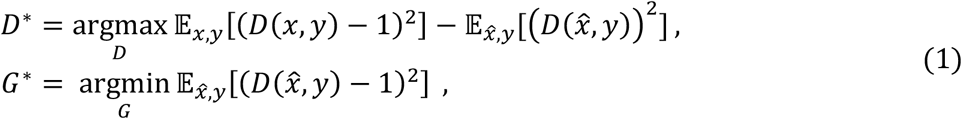

where 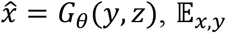 is the expectation over the true joint probability distribution of (*X, y*), and 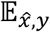is the joint probability distribution represented by the generator *G*. For additional details, we direct interested readers to (Pyne et al., 2026).

Once DiViA-mGAN is trained, the trained generator *G*^*^ can be used to generate trajectories for a new participant with measurements 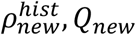 via the following procedure (also see Phase 3 in Figure 1):

1. Sample *Z*~*pz* and construct 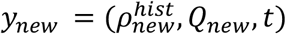 for a future time point *t*
2. Feed *Z, y*_*new*_ to *G*^*^ and obtain a possible value 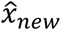 of the phenotype at time *t*.
3. For the fixed *Z*, vary the time *t* to generate a single plausible trajectory for the participant.
4. Repeat steps 1-3 by resampling *Z*~*pz* to generate other plausible trajectories.

The proposed DyViA-mGAN is an improvement over DyViA-GAN in that the generator network is constructed to model mixture effects. In particular, the generator has the form

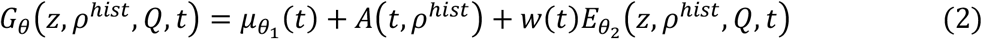

where 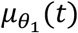 is a subnetwork (with trainable parameters *θ*_1_) modeling the population level dynamics, 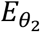 is a subnetwork (with trainable parameters *θ*_2_) modeling the subject specific dynamics, and *A*(*t, ρ*^*hist*^) is a correction term that ensures the generator’s predictions always satisfy the available phenotype history *ρ*^*hist*^ at the associated time points. Finally, *w(t)* is a polynomial weighting function that vanishes at the time points associated with *ρ*^*hist*^, i.e., *w*(*t*_*i*_) = 0 for *i* = 1, …, *N*. The weighting function is essential for the corrector term *A* to work as intended.

Additional details about the network architecture and hyperparameters can be found in the Supplementary Material.

## DyViA-Diff

Our main contribution is the design of a cSDM named DyViA-Diff. The underlying principle of a cSDM is that that the target conditional distribution *px*_|*y*_(*x*|*y*) can be transformed into a Gaussian distribution using a Fokker-Plank equation, which is a partial differential equation solving for the evolving distribution *p*^*τ*^(*x*|*y*). Here *τ* is a pseudo-time with *p*^0^(*x*|*y*) = *px*_|*y*_(*x*|*y*) and *p*^*T*^(*x*|*y*) = *N*(0, σ^2^(*T*)*I*) at some final time *τ* = *T*, where σ^2^(*τ*) is the noise variance schedule. At the level of individual samples, a sample from *x*_0_ ~ *p*^0^(*x*|*y*) is transformed into a sample *x*_*T*_ from the Gaussian *p*^*T*^(*x*|*y*) by sequentially injecting noise. We can then solve the “reverse” denoising process of transforming a sample *x*_*T*_ ~ *p*^*T*^(*x*|*y*) into a target sample *x*_0_ ~ *p*^0^(*x*|*y*), with the denoising governed by an Itô stochastic differential equation (see Supplementary Material for details).

Solving the reverse process requires access to the *score function s*(*x, τ, y*) = ∇_*x*_ log *p*^*τ*^(*x*|*y*), which provides the appropriate direction of denoising to move closer to realistic samples from *p*^0^(*x*|*y*) = *px*_|*y*_(*x*|*y*). This score function is approximated using a neural network *s*_*θ*_(*x, τ, y*) called the *score network* (with trainable parameters *θ*) by minimizing the following score matching objective

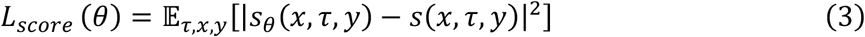

Via a clever manipulation, the objective (3) can be written (approximately) in terms of the labelled data D. Further details about the training process can be found in (Dasgupta et al., 2026; Tashiro et al., 2021).

Using the trained score network, which we denote as *s*^*^(*x, τ, y*) the stochastic differential equation associated with the denoising process can be solved (discretely) to generate the required samples from *px*_|*y*_(*x*|*y*). In particular, we can generate phenotype trajectories for a new participant with measurements *ρ*^*hist*^, *Q*_new_ via the following procedure (also see Phase 3 in Figure 1):

1. Sample *x*_*T*_~*N*(0, *I*) and construct 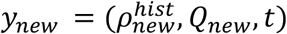 for a future time point *t*
2. Using the score function *s*^*^, solve the reverse denoising process to obtain a possible value 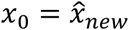 of the phenotype at time *t*.
3. For the fixed *x*_*T*_, vary the time *t* to generate a single plausible trajectory for the participant.
4. Repeat steps 1-3 by resampling *x*_*T*_~*N*(0, *I*) to generate other plausible trajectories.

Note the similarity of the above procedure with that of DyViA-mGAN. Additional details about the score network architecture and DyViA-Diff hyperparameters can be found in the Supplementary Material.

## Dataset

We modeled the anonymized longitudinal data from the *Study of Osteoporotic Fracture (SOF)*, which was a nationwide multicenter research study that began in 1986, funded by the U.S. National Institutes of Health (NIH) and is available publicly. The study enrolled 9,704 primarily Caucasian women and continued to track them over follow-up clinical visits. The dataset is publicly available from the ‘SOF Online’ website^2^. As proof of concept, we choose the phenotype of interest to be the femoral neck BMD (denoted by FND) measured by DXA scans, and computed the corresponding T-scores (*ρ*) for trajectory generation. Our data processing steps are described in the Supplementary Material.

We use data from four clinical visits, denoting the participant ages at the visits by *t*_1_, *t*_2_, *t*_3_, *t*_4_ and the corresponding FND values as *ρ*(*t*_1_), *ρ*(*t*_2_), *ρ*(*t*_3_), *ρ*(*t*_4_). The data used to construct *ρ*^*hist*^ for each participant is based on the measurements at *t*_1_, *t*_2_, i.e., *K* = 2. The remaining two temporal measurements are used to assess the accuracy of future predictions for test participants.

## Results

With each of the proposed generative models, we generate *L* = 1000 personalized plausible trajectories of the FND evolution for each test sample. These can then be used to sample mean 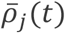 for each test participant *j*, conditioned on the known measurement *y*_*j*_. This mean is taken single trajectory reconstruction capturing the predicted personalized future dynamics of an individual’s phenotype.

### Performance metrics

We assess the accuracy of predictions using two metrics

- *Root mean square error (RMSE):* For each test participant *j*, we compute the squared error between 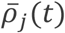 and the true values *ρ*_*j*_*(t)* at the two future time points *t*_3_, *t*_4_

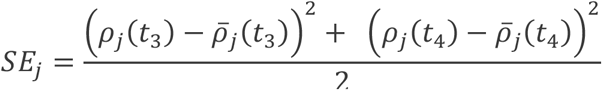

which is then used to compute the test RMSE as

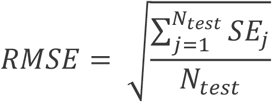 This serves as a quantitative metric to assess the prediction accuracy.
- *Mean and sample trajectories:* A qualitative assessment of the prediction is made by visualizing sample trajectories for four test participants. In addition, we plot the mean trajectory 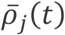 along with the 90% credible interval bounds at each time point along the trajectory (the 5^th^ and 95^th^ percentiles at each *t*). In each case, these are compared to the true values at *t*_1_, *t*_2_ (used for conditioning the predictions) and *t*_3_, *t*_4_.

### Model comparison

For DyViA-mGAN the test RMSE is 0.4411, while for DyViA-Diff this error is lower at 0.4364. Thus DyViA-Diff performs quantitative better than the cGAN-based model. The individual trajectories (10 per test participant) and the mean trajectories with (shaded) 90% credible interval bounds area shown in Figures 2 and 3, respectively. We immediately notice that:

**Figure 2:**
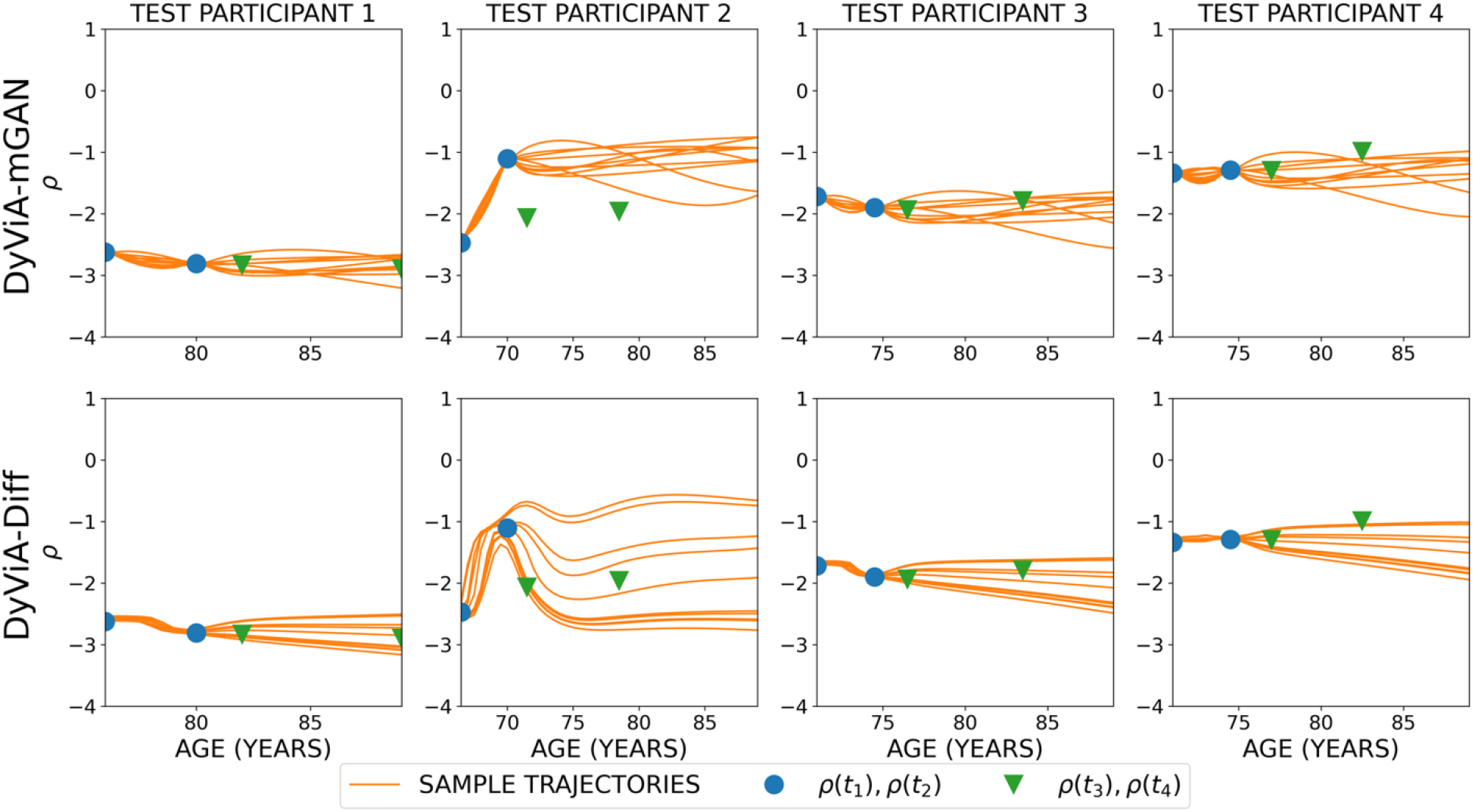
Comparing the quality of sample trajectories generated by DyViA-mGAN (top row) and DyViA-Diff (bottom row) for four test participants. 10 sample trajectories are shown for each participant.

**Figure 3:**
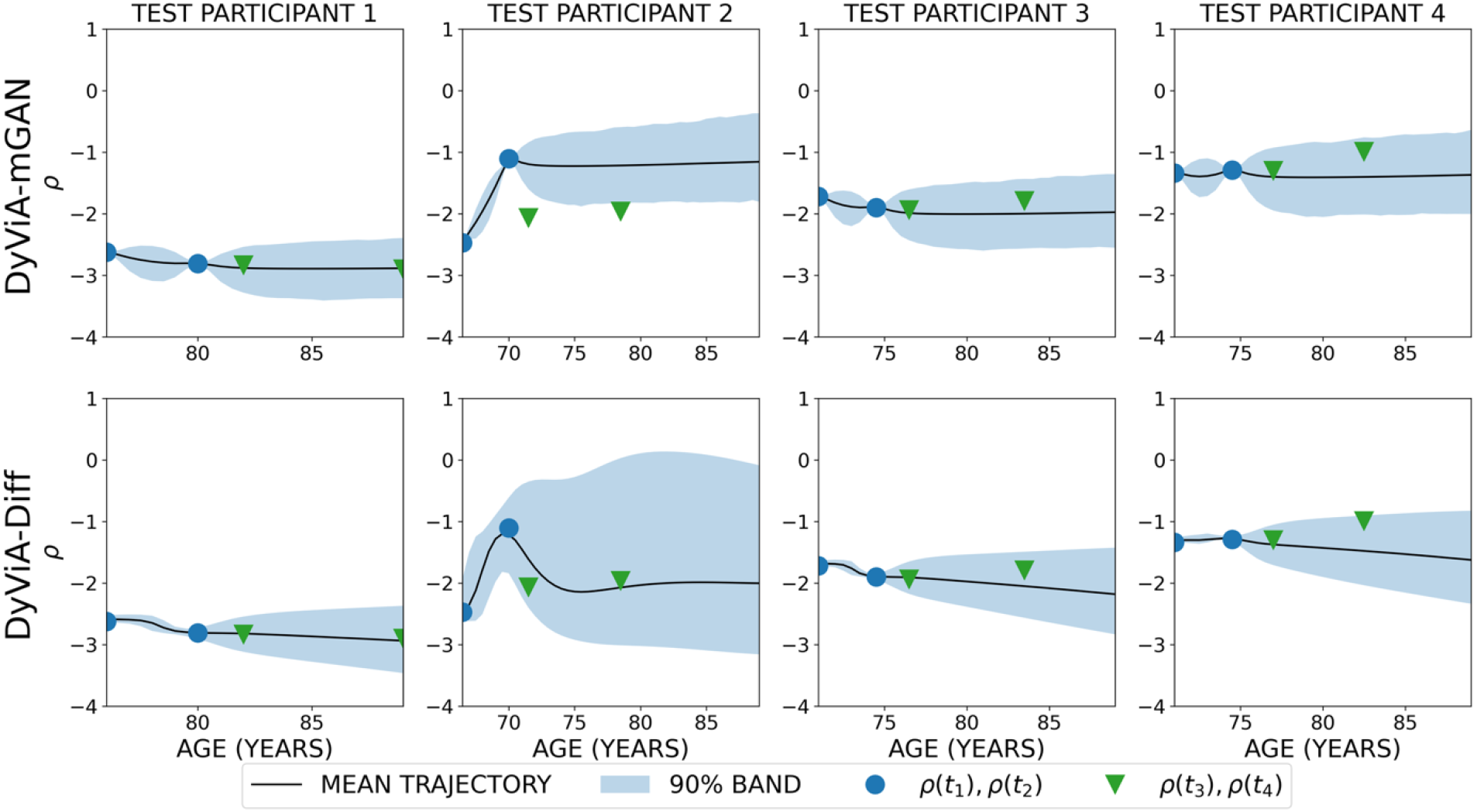
Comparing the quality of mean trajectories generated by DyViA-mGAN (top row) and DyViA-Diff (bottom row) for four test participants. The mean is taken of 1000 sample trajectories for each participant, which 90% credible intervals shown as shaded bands.

- Despite not having built-in corrector terms (as is done for DyViA-mGAN in (2)) to guarantee that the predicted samples match the known conditioning history *ρ*^*hist*^ (shown as blue dots), the sample trajectories with DyViA-Diff are reasonably consistent with *ρ*^*hist*^.
- When compared to DyViA-mGAN, the true values at future time points *t*_3_, *t*_4_ are better encapsulated by 90% credible intervals of DyViA-Diff while also ensuring tighter variability closer to the condition time points *t*_1_, *t*_2_ (narrower shaded region closer to known initial measurements in Figure 3).

The supplementary material shows additional comparisons with a standard LMM and the original DyViA-GAN (Pyne et al., 2026), and explores a filtering strategy to remove outlier trajectories. From an innovation perspective, while the original DyViA-GAN model outperforms standard statistical models such as the LMM, the new DyViA-mGAN enables us to encode population- and individual-level effects with a better control on the predicted trajectories (see Supplementary Materials). Notably, the proposed DyViA-Diff is the superior model both in terms of predicting coherent personalized trajectories as well as accuracy.

## Discussion and implications

It was estimated by a study of BMD that 43.4 and 10.2 million U.S. adults of age 50 years and above have low bone mass and osteoporosis, respectively (Wright et al., 2014). One osteoporosis study reported that, given the anticipated population aging and growth in the U.S., the annual projection figures could escalate up to 3.2 million fractures with related costs of over $95 billion by 2040 (Lewiecki et al., 2019). It underscores the urgent need for evidence-based and cost-effective anti-osteoporosis treatments (Levis & Theodore, 2012) and possible lifestyle strategies (Potashkin & Kim, 2024). Individuals often lose about 0.5-1% of BMD each year after the age of 50, with an even sharper decline of 1-3% for women 5-10 years post-menopause. Preventive interventions on the one hand and histories of fractures and certain diseases on the other could influence individual BMD trajectories, e.g., (Rathbun et al., 2016). Of emerging importance are the varied effects of weight-loss interventions on bone health, a topic of current investigation (Paccou et al., 2025).

The main impact of DyViA’s genAI-driven contribution in the form of multiple personalized, plausible models of BMD progression with age lies in the systematic yet rigorous transition from otherwise static models of inference about a clearly dynamic phenomenon to a continuous one. DyViA does this with healthy BMD trajectories and without any disruption of the upstream densitometric data analysis or downstream assessment of fracture risks and clinical diagnosis of osteoporosis. From a practical perspective, the foresight of such a plausible range of trajectories can empower an individual by conferring a certain degree of strategic preparedness in the course of aging, e.g., lifestyle modifications, optimal built environment, effective insurance and financial planning.

As its genAI models become more powerful, DyViA’s output could potentially be used as an aid in clinical decision-making as well as preventive and personalized medicine. Indeed, its flexible suite of algorithms could allow the trajectories in the future to be dynamically updated through modifiable parameters that inform the DyViA models of possible individual-specific intermediate events including fractures, lifestyle alterations, or medical interventions. Meanwhile, the authors will work on addressing current limitations of the platform such as training of the models with further heterogeneous longitudinal data from various populations and groups as well as other relevant phenotypes of aging. Available data on chronic comorbidities and common risk factors of osteoporosis as well as its secondary contributors including certain medication use could be considered for individual-specific stratification and prediction by the future versions of our genAI models.

## Footnotes

1 *N*(*µ*, σ^2^*I*) denotes a multivariate uncorrelated Gaussian with mean *µ* and diagonal covariance σ^2^*I* with σ^2^ > 0.

2 https://sofonline.ucsf.edu/

